# Large scale genome-wide association study in a Japanese population identified 45 novel susceptibility loci for 22 diseases

**DOI:** 10.1101/795948

**Authors:** Kazuyoshi Ishigaki, Masato Akiyama, Masahiro Kanai, Atsushi Takahashi, Eiryo Kawakami, Hiroki Sugishita, Saori Sakaue, Nana Matoba, Siew-Kee Low, Yukinori Okada, Chikashi Terao, Tiffany Amariuta, Steven Gazal, Yuta Kochi, Momoko Horikoshi, Ken Suzuki, Kaoru Ito, Yukihide Momozawa, Makoto Hirata, Koichi Matsuda, Masashi Ikeda, Nakao Iwata, Shiro Ikegawa, Ikuyo Kou, Toshihiro Tanaka, Hidewaki Nakagawa, Akari Suzuki, Tomomitsu Hirota, Mayumi Tamari, Kazuaki Chayama, Daiki Miki, Masaki Mori, Satoshi Nagayama, Yataro Daigo, Yoshio Miki, Toyomasa Katagiri, Osamu Ogawa, Wataru Obara, Hidemi Ito, Teruhiko Yoshida, Issei Imoto, Takashi Takahashi, Chizu Tanikawa, Takao Suzuki, Nobuaki Sinozaki, Shiro Minami, Hiroki Yamaguchi, Satoshi Asai, Yasuo Takahashi, Ken Yamaji, Kazuhisa Takahashi, Tomoaki Fujioka, Ryo Takata, Hideki Yanai, Akihide Masumoto, Yukihiro Koretsune, Hiromu Kutsumi, Masahiko Higashiyama, Shigeo Murayama, Naoko Minegishi, Kichiya Suzuki, Kozo Tanno, Atsushi Shimizu, Taiki Yamaji, Motoki Iwasaki, Norie Sawada, Hirokazu Uemura, Keitaro Tanaka, Mariko Naito, Makoto Sasaki, Kenji Wakai, Shoichiro Tsugane, Masayuki Yamamoto, Kazuhiko Yamamoto, Yoshinori Murakami, Yusuke Nakamura, Soumya Raychaudhuri, Johji Inazawa, Toshimasa Yamauchi, Takashi Kadowaki, Michiaki Kubo, Yoichiro Kamatani

**Author notes:** Correspondence and requests for materials should be addressed to: Soumya Raychaudhuri, M.D., Ph.D., Center for Data Sciences, Harvard Medical School, Boston, MA, USA. Yoichiro Kamatani, M.D., Ph.D., Laboratory for Statistical Analysis, RIKEN Center for Integrative Medical Sciences, Yokohama, Japan.

## Abstract

The overwhelming majority of participants in current genetic studies are of European ancestry^1–3^, limiting our genetic understanding of complex disease in non-European populations. To address this, we aimed to elucidate polygenic disease biology in the East Asian population by conducting a genome-wide association study (GWAS) with 212,453 Japanese individuals across 42 diseases. We detected 383 independent signals in 331 loci for 30 diseases, among which 45 loci were novel (*P* < 5 × 10^−8^). Compared with known variants, novel variants have lower frequency in European populations but comparable frequency in East Asian populations, suggesting the advantage of this study in discovering these novel variants. Three novel signals were in linkage disequilibrium (r^2^ > 0.6) with missense variants which are monomorphic in European populations (1000 Genomes Project) including rs11235604(p.R220W of *ATG16L2*, a autophagy-related gene) associated with coronary artery disease. We further investigated enrichment of heritability within 2,868 annotations of genome-wide transcription factor occupancy, andidentified 378 significant enrichments across nine diseases (FDR < 0.05) (e.g. NF-κB for immune-related diseases). This large-scale GWAS in a Japanese population provides insights into the etiology of common complex diseases and highlights the importance of performing GWAS in non-European populations.

## MAIN TEXT

We conducted a genome-wide association study (GWAS) of 42 diseases in a Japanese population, comprising 179,660 patients who participated in the BioBank Japan Project (BBJ)^4,5^ and 32,793 population-based controls (Supplementary Table 1). The 42 diseases encompassed a wide-range of disease categories; 13 neoplastic diseases, five cardiovascular diseases, four allergic diseases, three infectious diseases, two autoimmune diseases, one metabolic disease, and 14 uncategorized diseases. By including patients with unrelated diagnoses into control samples, we maximized the power of our GWAS (Supplementary Table 1 and Supplementary Figure 1). We employed a generalized linear mixed model in our association analysis using SAIGE^6^. Following imputation with 1000 Genomes Project Phase 3 reference data (1KG Phase3)^7^, we tested 8,712,794 autosomal variants and 207,198 X chromosome variants for association with 42 diseases. We estimated their heritability using linkage disequilibrium score regression (LDSC) analysis^8^ (Supplementary Table 2). Consistent with a recent finding in the European population^9^, incorporating the baselineLD model^10^ into the LDSC analysis improved heritability estimation in our GWAS (Methods; Supplementary Figure 2 and Supplementary Table 2). Although we observed high genomic inflation factors (*λ*_*GC*_) for some diseases (e.g. *λ*_*GC*_ = 1.3 for type 2 diabetes (T2D); Supplementary Table 2), LDSC analysis indicated that the majority of the inflated chi-squared statistics originated from polygenic effects rather than confounding biases (e.g. intercept = 1.01 for T2D; Supplementary Table 2). Overall, we detected significant associations for 30 diseases at 309 autosomal loci (outside of the HLA region) and nine loci on the X chromosome (*P* < 5 × 10^−8^) (Supplementary Table 3 and 4). Associations at the HLA region have been investigated in detail in a separate article^11^.

**Table 1.**
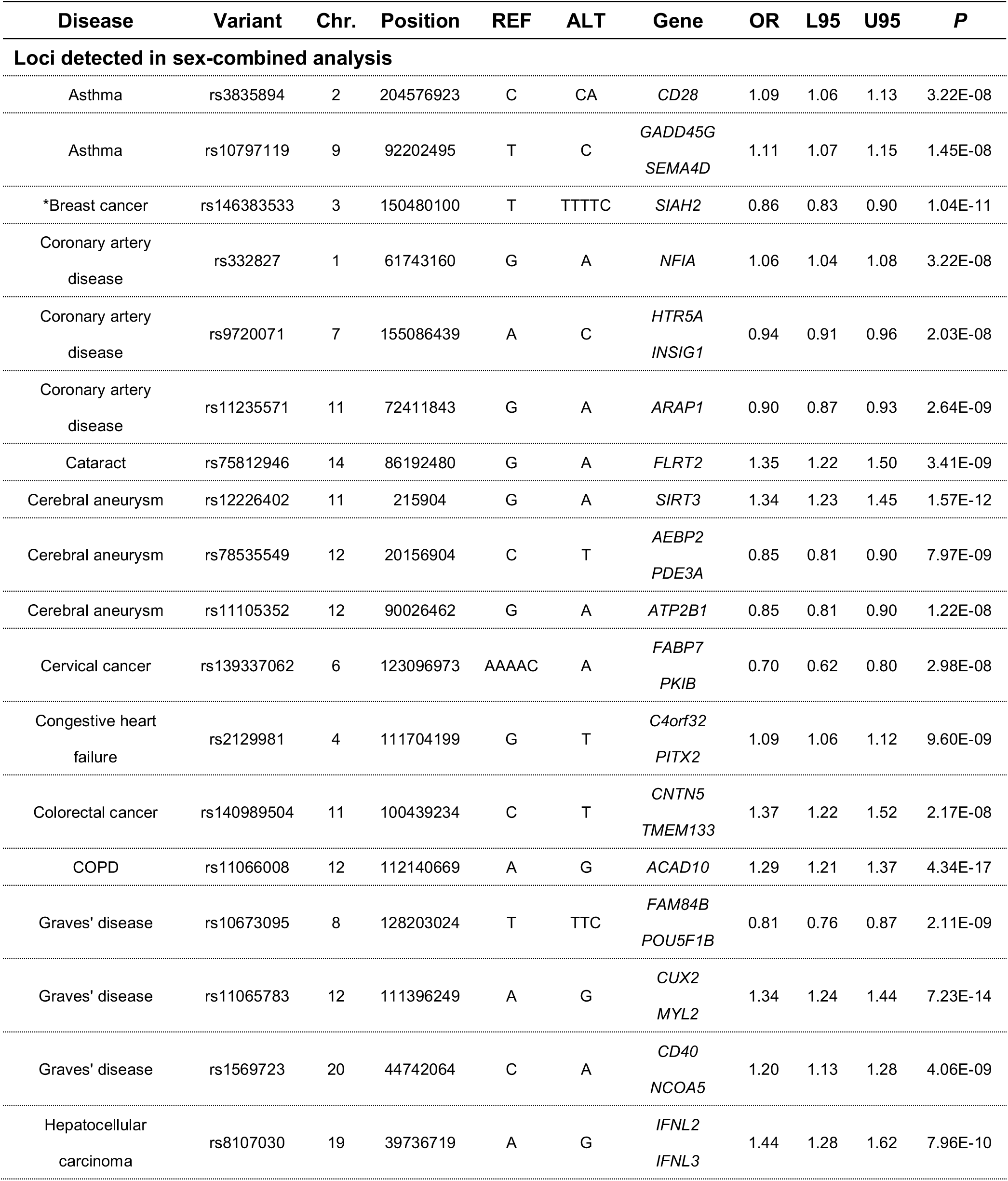
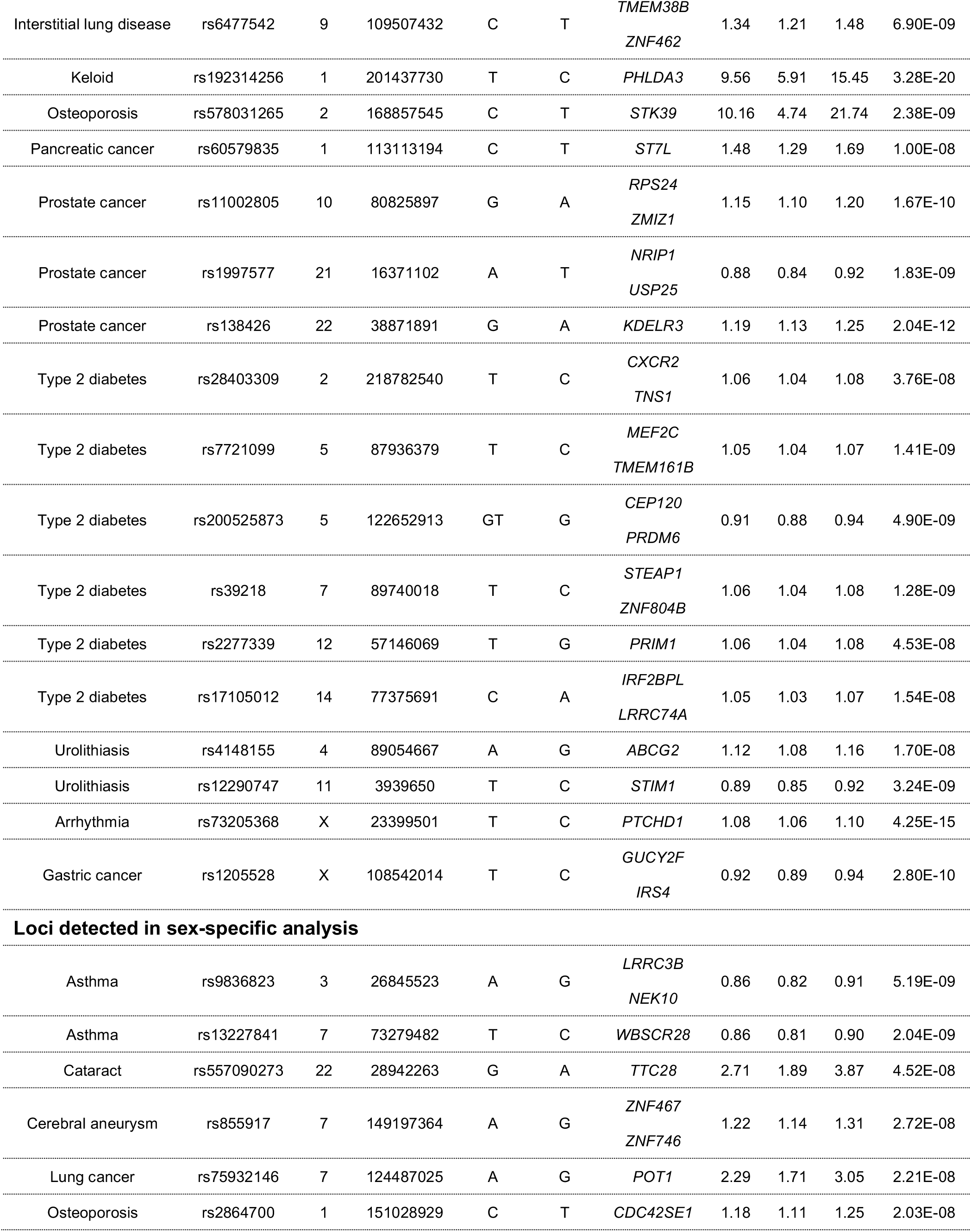
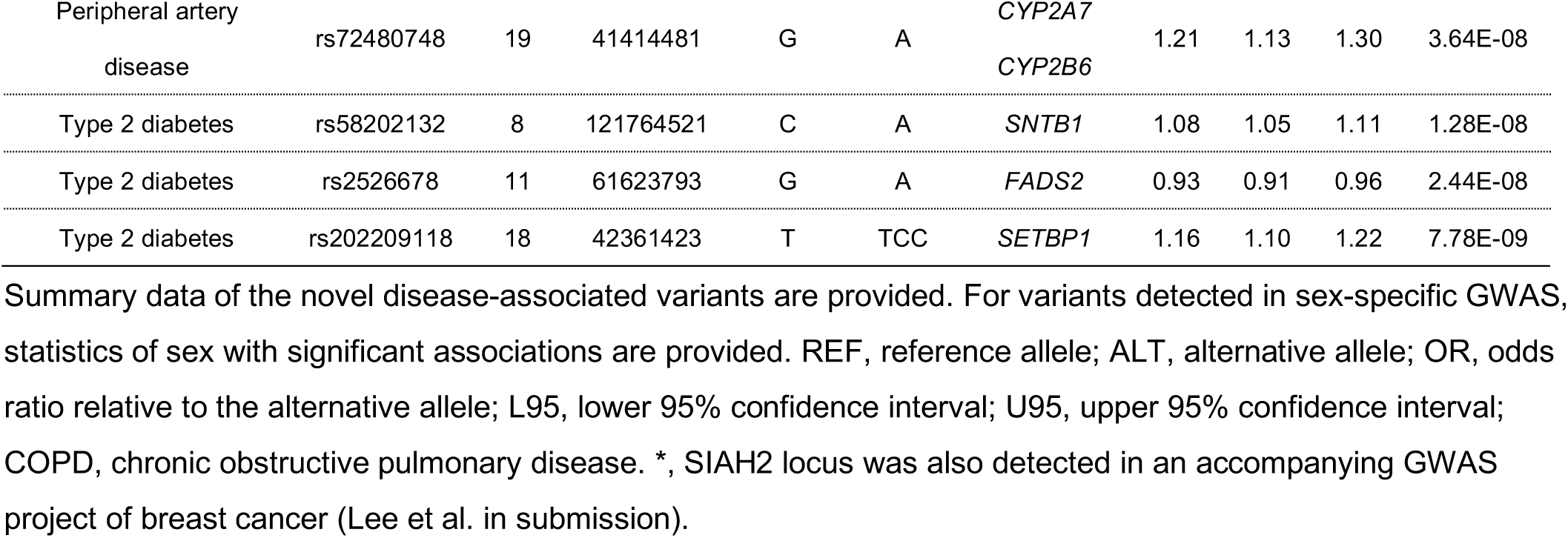
45 novel loci detected in this GWAS.

**Table 2.** Missense variants in LD with nine novel disease-associated variants.

|  |  |  |  |  |  |  |  |  |  |  |  | Allele freq. in<br>1KG Phase3 |  |  |
| --- | --- | --- | --- | --- | --- | --- | --- | --- | --- | --- | --- | --- | --- | --- |
| Disease | Variant | Chr. | Position | Gene | Amino<br>acid<br>change | REF | ALT | OR | L95 | U95 | P | EAS | EUR | AFR |
| Loci detected in sex-combined analysis |  |  |  |  |  |  |  |  |  |  |  |  |  |  |
| Coronary<br>artery disease | rs11235604 | 11 | 72533536 | ATG16L2 | p.R220W | C | T | 0.91 | 0.88 | 0.94 | 1.73E-08 | 0.100 | 0.000 | 0.000 |
| Hepatocellular<br>carcinoma* | rs8103142 | 19 | 39735106 | IFNL3 | p.K70R | T | C | 1.38 | 1.23 | 1.54 | 1.14E-08 | 0.083 | 0.312 | 0.695 |
| Keloid | rs192314256 | 1 | 201437730 | PHLDA3 | p.E62G | T | C | 9.56 | 5.91 | 15.45 | 3.28E-20 | 0.010 | 0.000 | 0.000 |
| Type 2 diabetes | rs2303720 | 5 | 122682334 | CEP120 | p.R921H<br>p.R947H | C | T | 0.91 | 0.89 | 0.94 | 1.77E-08 | 0.084 | 0.026 | 0.048 |
| Type 2 diabetes | rs194520 | 7 | 89854446 | STEAP2 | p.F17C | T | G | 1.06 | 1.04 | 1.08 | 1.23E-08 | 0.183 | 0.512 | 0.286 |
| Type 2 diabetes | rs194524 | 7 | 89861832 | STEAP2 | p.R456Q | G | A | 1.06 | 1.04 | 1.08 | 1.18E-08 | 0.183 | 0.512 | 0.269 |
| Type 2 diabetes | rs2277339 | 12 | 57146069 | PRIM1 | p.D5A | T | G | 1.06 | 1.04 | 1.08 | 4.53E-08 | 0.206 | 0.111 | 0.199 |
| Urolithiasis | rs2231142 | 4 | 89052323 | ABCG2 | p.Q141K | G | T | 1.12 | 1.08 | 1.16 | 1.74E-08 | 0.291 | 0.094 | 0.013 |
| Loci detected in sex-specific analysis |  |  |  |  |  |  |  |  |  |  |  |  |  |  |
| Asthma | rs13246460 | 7 | 73249165 | WBSCR27 | p.R216W | T | A | 0.89 | 0.85 | 0.93 | 9.23E-07 | 0.581 | 0.726 | 0.463 |
| Asthma | rs13232463 | 7 | 73249299 | WBSCR27 | p.S171W | G | C | 0.89 | 0.85 | 0.93 | 9.23E-07 | 0.581 | 0.726 | 0.463 |
| Asthma | rs13241921 | 7 | 73254812 | WBSCR27 | p.Q107R | T | C | 0.89 | 0.85 | 0.93 | 9.54E-07 | 0.581 | 0.726 | 0.464 |
| Asthma | rs11770052 | 7 | 73275565 | WBSCR28 | p.I14N | T | A | 0.87 | 0.83 | 0.92 | 3.20E-08 | 0.630 | 0.726 | 0.285 |
| Asthma | rs13227841 | 7 | 73279482 | WBSCR28 | p.W78R | T | C | 0.86 | 0.81 | 0.90 | 2.04E-09 | 0.650 | 0.677 | 0.334 |
| Lung cancer | rs75932146 | 7 | 124487025 | POT1 | p.V326A | A | G | 2.29 | 1.71 | 3.05 | 2.21E-08 | 0.003 | 0.000 | 0.000 |

Summary data of the missense variants in LD ( $r^2 > 0.6$ ) with novel disease-associated variants are provided. For variants detected in sex-specific GWAS, statistics of sex with significant associations are provided. Rows highlighted in bold indicate east Asian-specific variants (MAF = 0 in Europeans of 1KG Phase3). REF, reference allele; ALT, alternative allele; Allele freq., allele frequency of alternative allele; OR, odds ratio relative to the alternative allele; L95, lower 95% confidence interval; U95, upper 95% confidence interval; EAS, East Asian populations; EUR, European populations; and AFR, African populations. \*, the lead variant at this region (rs8107030) is in LD ( $r^2 = 0.880$ ) with ss469415590 (comprised of rs67272382 and rs74597329). ss469415590 is a frameshift variant of *IFNL4*<sup>37</sup> (not listed as a functional gene in GRCh37 coordinates), and it might also explain this association.

**Figure 1.**
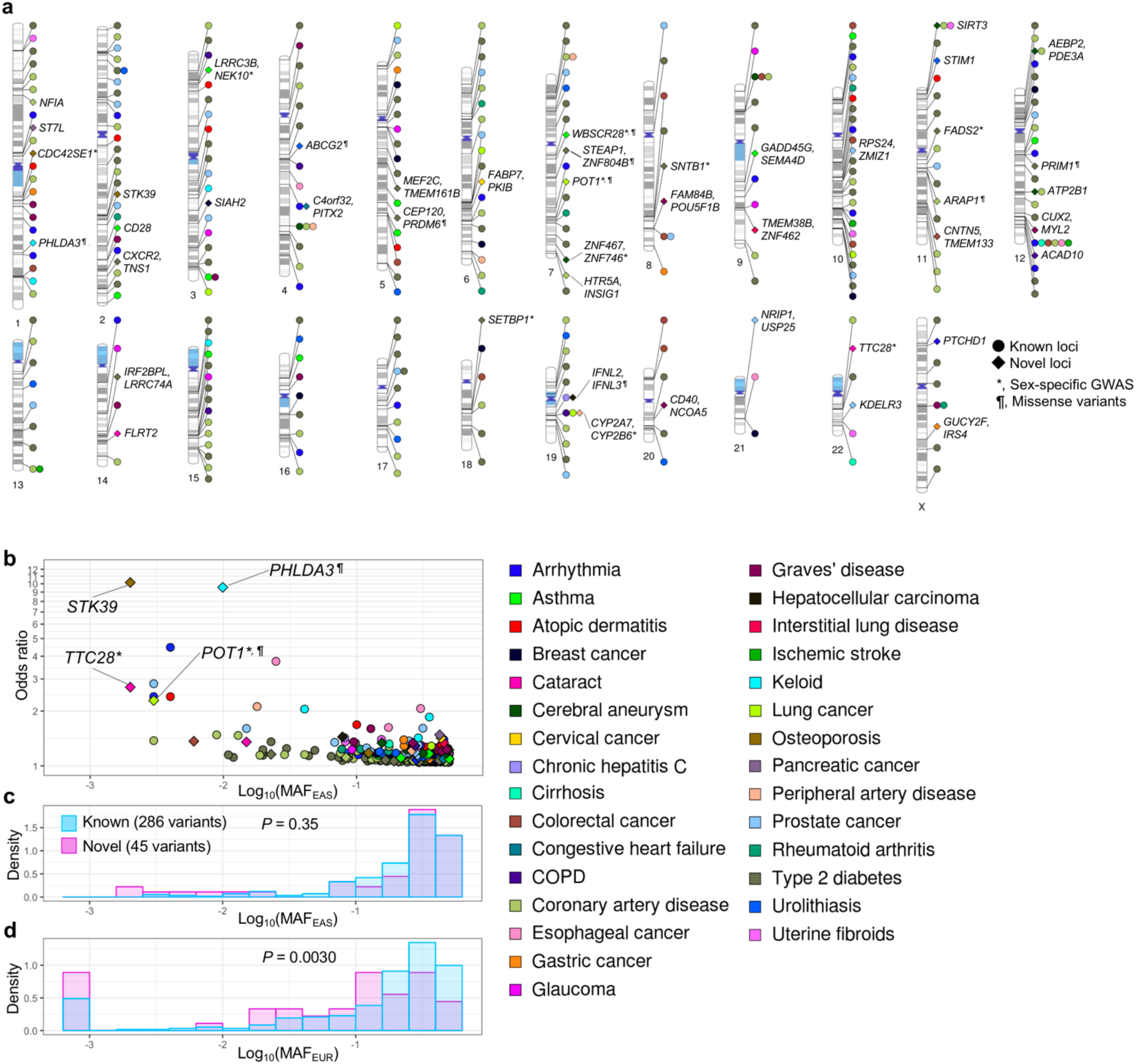
Overview of 331 loci detected in this GWAS. **a**, Phenogram^38^ of 331 loci detected in this GWAS. Novel loci (♦) were annotated by the closest gene names. Pleiotropic associations (see Methods for its definition) were plotted at the same position. **b**, Allele frequencies and the odds ratios of the lead variants at 331 loci detected in this GWAS. The odds ratio of the risk allele was used. *, loci detected in sex-specific GWAS. ¶, the lead variants are in LD with missense variants (r^2^ > 0.6). **c**, **d**, Allele frequencies of the lead variants were compared between novel loci and known loci (**c**, East Asian populations; **d**, European populations). The difference in MAF was tested by Mann–Whitney U test, and its *P* value was provided. When MAF < 0.001, MAF was adjusted to 0.001 to fit in log scale. MAF_EAS_, MAF in East Asian population (1KG Phase3). MAF_EUR_, MAF in European population (1KG Phase3).

**Figure 2.**
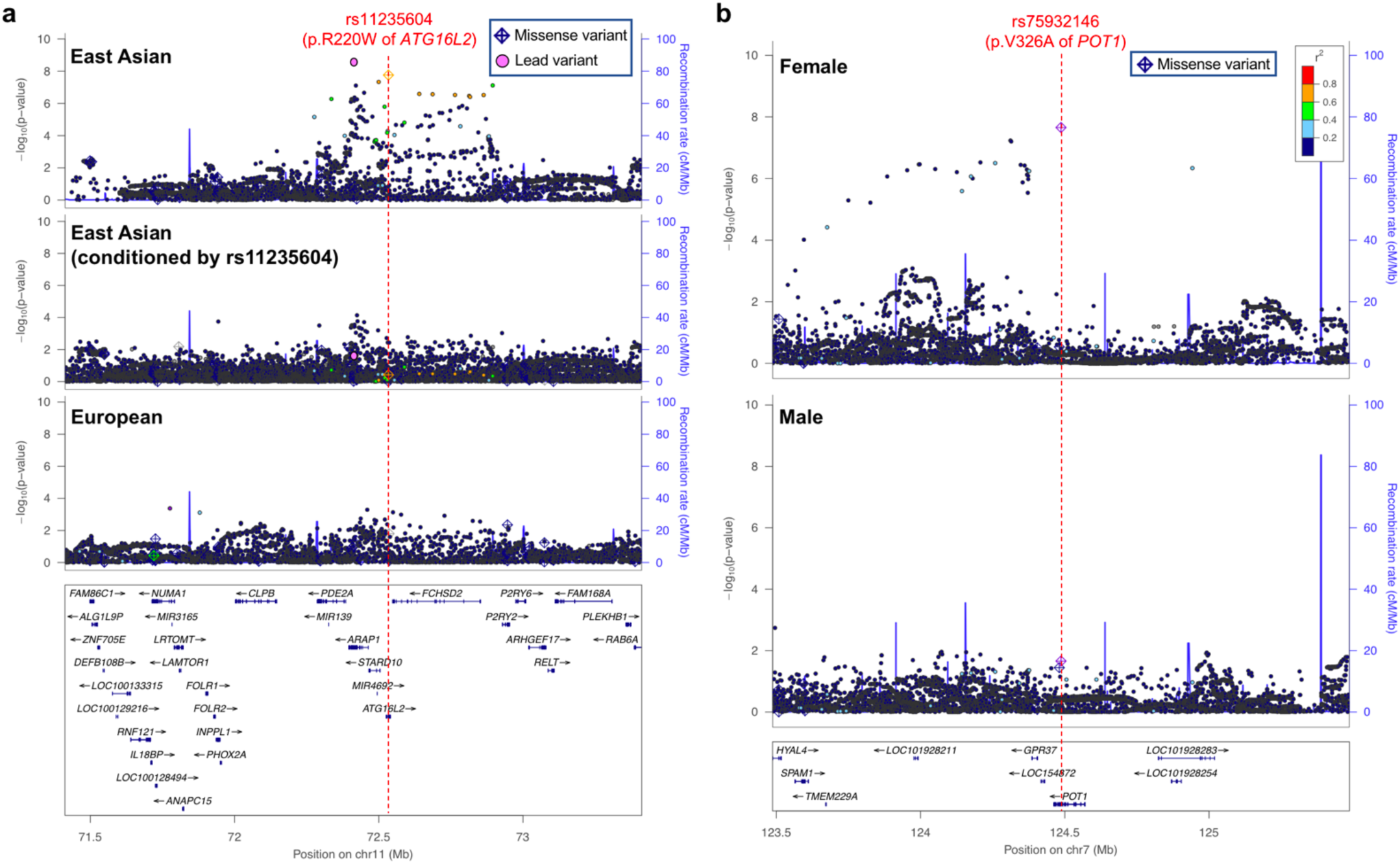
Novel associations which can be explained by East Asian-specific missense variants. Regional association plots are provided (**a**, coronary artery disease; and **b**, lung cancer). For coronary artery disease (**a**), *P* values from conditional analysis and those in European GWAS^12^ were plotted separately. For lung cancer (**b**), *P* values from female- and male-specific GWAS were plotted separately.

We further performed conditional analyses in these 318 loci to explore disease-associated variants independent of the lead variants. We detected 52 additional independent signals for 11 diseases (*P* < 5 × 10^−8^) (Supplementary Table 5). The largest number of independent signals in a single locus was seven, found in the *FAM84B/POU5F1B* locus associated with prostate cancer and in the *KCNQ1* locus associated with T2D.

For 35 diseases for which we have both male and female patients, we conducted male- and female-specific GWAS. We detected 13 additional loci for 10 diseases which were not identified in a sex-combined analysis (*P* < 5 × 10^−8^) (Supplementary Table 6). We tested heterogeneity between effect size estimates for males and females using Cochran’s Q test. This analysis found seven loci showing significant differences in effect size estimates between sexes (*P* values of heterogeneity (*P*_*het*_) < 0.05/13); two asthma loci, a cataract locus, a cerebral aneurysm locus, and a lung cancer locus were specifically associated with females; and a coronary artery disease (CAD) locus and a T2D locus were specifically associated with males.

In total, we detected 383 independent signals in 331 loci outside of the HLA region for 30 diseases, of which 45 loci were novel (Figure 1a, Table 1, Supplementary Table 3, 4, and 6). Five novel disease-associated variants were rare variants (MAF < 0.01), and four of them had large effect sizes (odds ratio > 2, Figure 1b). To understand the characteristics of novel and known disease-associated variants, we examined their allele frequencies in East Asian and European populations of 1KG Phase3. Allele frequencies of novel and known variants were of comparable level in East Asian populations (*P* = 0.35, Figure 1c). However, novel variants have lower allele frequencies than known variants in European populations (*P* = 0.0030, Figure 1d). Although both of novel and known variants have lower allele frequencies in European populations than in East Asian populations, novel variants have larger inter-population differences than known variants (*P* = 0.0047, Supplementary Figure 3). To estimate population specificity in our GWAS results, we compared our results with those reported in previous European GWAS. We utilized publicly available GWAS summary statistics of European populations for 10 diseases (Methods), and tested for consistency in direction of effect between populations at 11 novel and 146 known disease-associated variants from our GWAS; 10 out of 11 novel and 141 out of 146 known variants were replicated in the same allelic direction in European GWAS (binomial test *P* values were 0.011 and 1.1 × 10^−35^, respectively; Supplementary Figure 4). In addition, 595 out of 665 disease-associated variants detected in European GWAS were replicated in the same allelic direction in our GWAS (binomial test *P* values = 1.1 × 10^−104^; Supplementary Figure 4). These findings suggested that genetic etiologies around the disease-associated variants are generally shared across populations, and the higher allele frequencies at novel associated variants in our East Asian cohort increased the efficiency of the variant discovery. This highlights the importance of performing GWAS in non-European populations.

**Figure 3.**
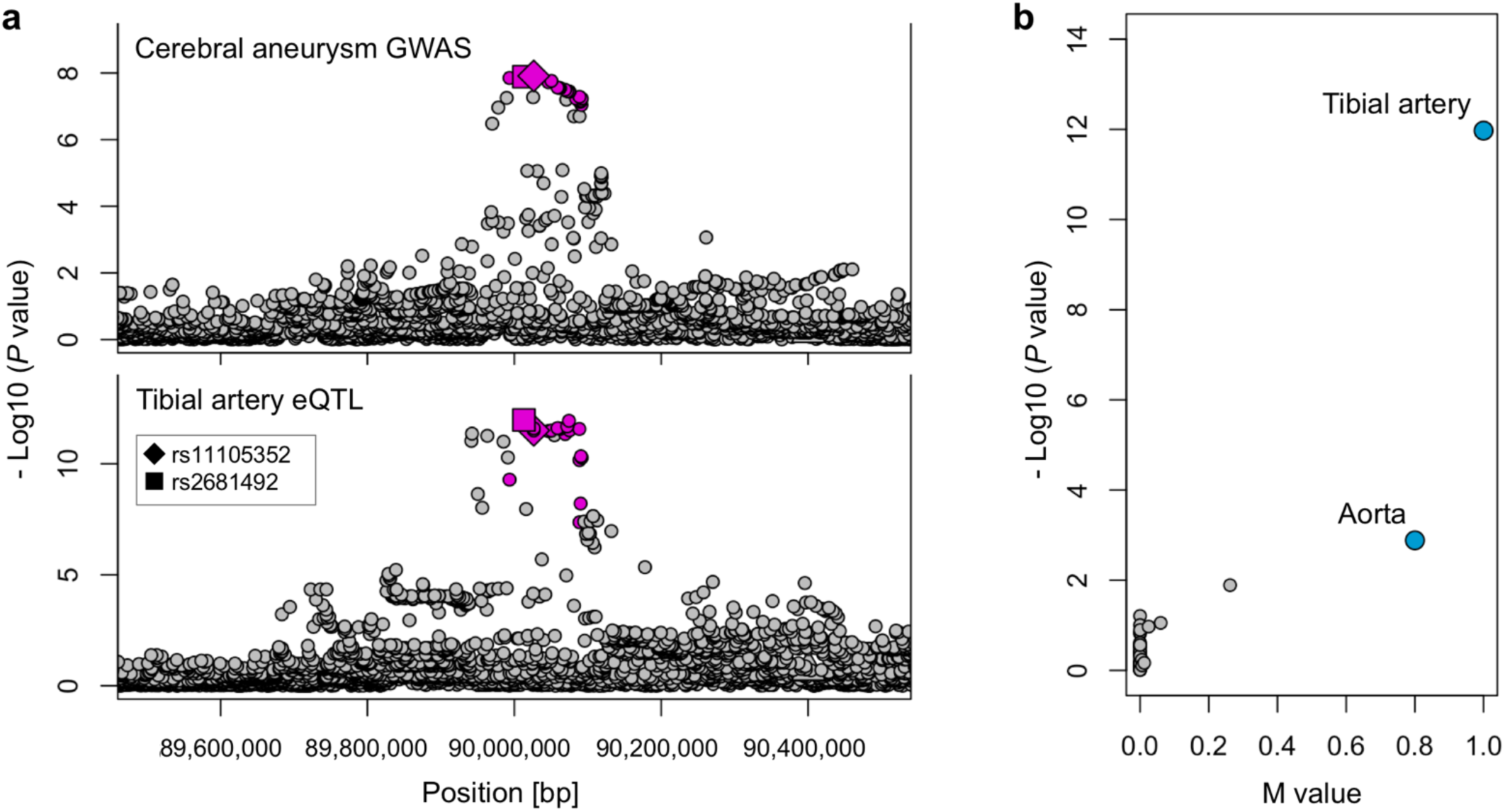
A novel association of cerebral aneurysm can be explained by artery-specific eQTL signals for *ATP2B1*. **a.** Regional association plots of cerebral aneurysm GWAS at *ATP2B1* locus (top) and those of eQTL signals for *ATP2B1* in the tibial artery (bottom) are provided. The lead variant of GWAS (rs11105352; ♦ dot) and the lead variant of eQTL (rs2681492; ▪ dot) are indicated by different shapes. Variants in LD with rs11105352 are highlighted by red (r^2^ > 0.6 both in East Asian and European populations of 1KG Phase3). **b**, Tissue-specificity of eQTL signals for *ATP2B1* at rs2681492 (the lead variant of eQTL in the tibial artery (▪ dot in **a**)). *P* values in eQTL analysis and M values (the posterior probability that an eQTL effect exist in each tissue tested in the cross-tissue meta-analysis) in all tissues in GTEx project^39^ are provided. Each dot indicates each tissue. All statistics of eQTL analysis were derived from release v7 of GTEx project^39^.

**Figure 4.**
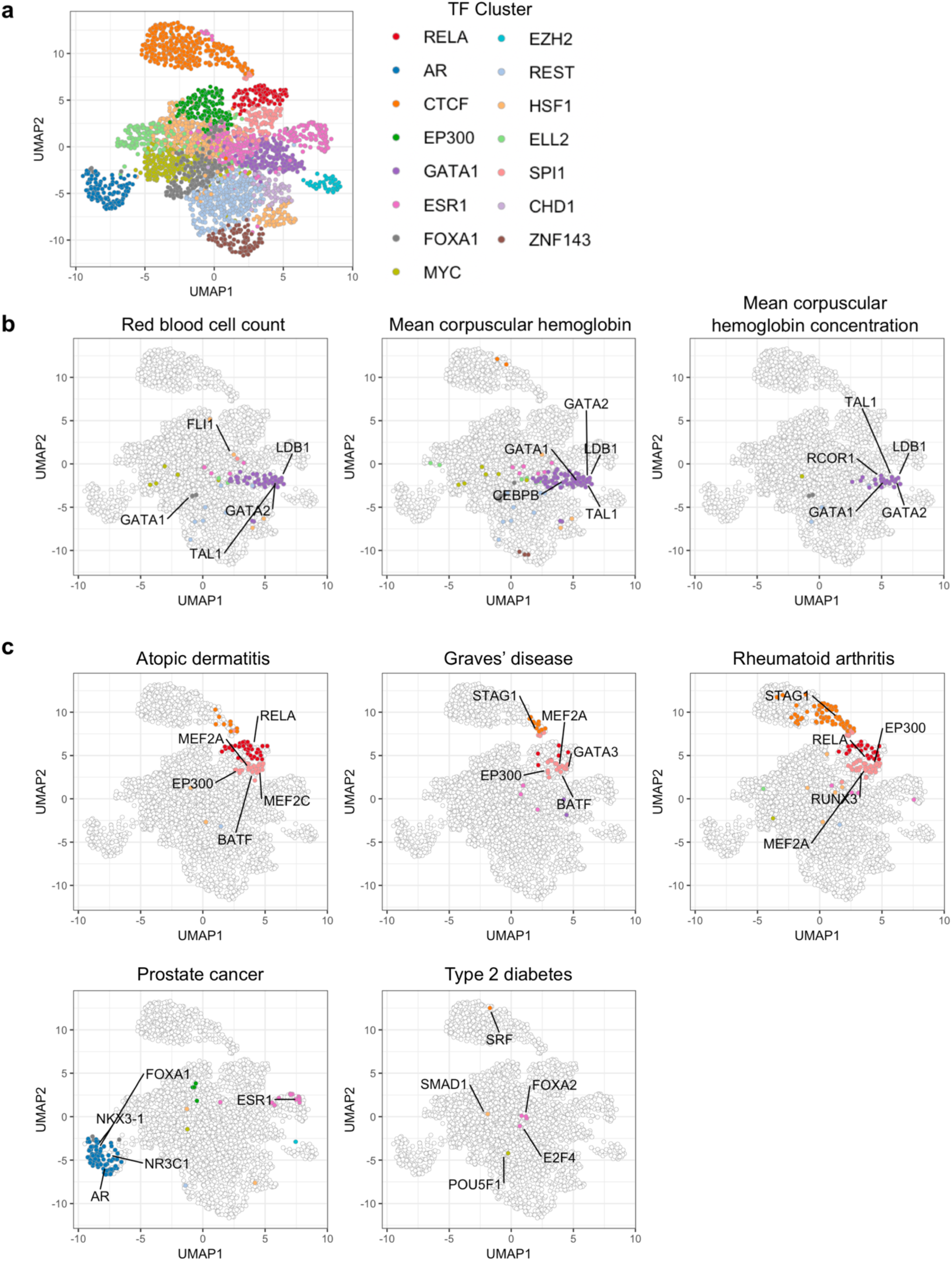
Transcription factors whose binding sites were enriched for heritability of diseases. **a**, All of the 2,868 sets of TF binding sites grouped into 15 clusters were plotted in the UMAP space. **b and c**, The results of S-LDSC were plotted on the UMAP space. The significant results (FDR < 0.05) are highlighted by cluster-specific colors. The names of the top five most significant TFs are also shown on the plot. **b**, The results of red blood cell-related traits. **c**, The results of diseases in this GWAS which had more than five significant TF binding site tracks (the results of other diseases are provided in Supplementary Figure 8).

We next investigated the potential impact of the disease-associated variants on protein functions (Supplementary Table 7). Nine novel variants were in linkage disequilibrium (LD) with missense variants (r^2^ > 0.6 in the 1KG Phase3 East Asian populations) (Table 2). Among them, three missense variants are monomorphic in European populations (1KG Phase3); p.R220W of *ATG16L2* associated with CAD; p.V326A of *POT1* associated with lung cancer; and p.E62G of *PHLDA3* associated with keloid (Figure 2 and Supplementary Figure 5). First, *ATG16L2* is an autophagy-related gene. Although p.R220W of *ATG16L2* is not the lead variant at this locus, conditioning on this variant cancelled the signal of the lead variant (Figure 2a). Previous GWAS for CAD in European populations did not detect significant associations at this locus^12^ (Figure 2a). These findings suggested that p.R220W of *ATG16L2* which is absent in Europeans may be the causal variant. p.R220W of *ATG16L2* is also associated with Crohn’s disease in a Chinese population^13^, and *ATG16L2* is highly expressed in immune cells. Therefore, dysregulated autophagy in immune cells might have an important role in CAD. Second, *POT1* is a member of the telombin family and this protein binds to telomeres, regulating telomere length. Missense variants of *POT1* have been described to be responsible for several familial cancers^14–16^. Together with a known association at the *TERT* locus (Supplementary Table 3), we provide additional evidence that telomere dysregulation is pathogenic for lung cancer. Intriguingly, this association was discovered in the female-specific GWAS, and a significant heterogeneity in effect sizes between males and females was observed (*P*_*het*_ = 7.7×810^−4^) (Figure 2b; Supplementary Table 6). This finding might help to understand the inter-sex differences in the etiology of lung cancer. Third, p.E62G of *PHLDA3* is predicted to have a deleterious effect to its protein function (SIFT score^17^=0; CADD score^18^=33), and we detected a large effect size for keloid (odds ratio = 9.56; 95% CI 5.91-15.45). PHLDA3 is known to be a suppressor of AKT^19^, and upregulated AKT signaling pathway is related to increased collagen production from dermal fibroblasts^20^. Therefore, damaged PHLDA3 may activate the AKT pathway, promoting the development of keloid. Together, our study successfully identified novel potential causal genes which would be hard to be discovered by GWAS in European populations due to restrictive European allele frequencies.

We also investigated the potential impacts of the disease-associated variants on the mRNA levels using two databases of expression quantitative trait locus (eQTL) analysis^21,22^. Out of the remaining 36 novel variants whose functions were not explained by missense variants,11 variants were in LD with at least one eQTL variant (r^2^ > 0.6), regulating the expression of 17 genes in total (Supplementary Table 8); *P2RY13, SIAH2*, and *SIAH2-AS1* for breast cancer; *ATP2B1, BET1L, POC1B*, and *ZNF767* for cerebral aneurysm; *CCAT1* and *CD40* for Graves’s disease; *GABPB2* for osteoporosis; *MOV10* and *WNT2B* for pancreatic cancer; *CYP2A6* and *CYP2B7P1* for peripheral artery disease; *ZMIZ1-AS1* for prostate cancer; and *STIM1* and *TRIM21* for urolithiasis. Intriguingly, the eQTL signals for *ATP2B1* which are in LD with a novel variant of cerebral aneurysm (rs11105352) is highly specific to arterial tissues (Figure 3). Since the loss of *ATP2B1* in vascular smooth muscle cells induced blood pressure elevation in mice^23^, decreased expression of *ATP2B1* in arteries might induce hypertension, which leads to increased risk of cerebral aneurysm.

To understand differences in the genetic risks between males and females, we assessed genetic correlations using LDSC^24^ between the results of sex-specific GWAS for the 20 diseases (see Methods for selection of diseases). Although most correlations are close to one, correlation of asthma was significantly smaller than one (*P* = 2.2 × 10^−3^ < 0.05/20; Supplementary Figure 6). This finding suggested that genetic risks of asthma is slightly different between males and females. To explore the biological mechanism underlying this finding, we estimated the enrichment of the heritability of male or female asthma in the 220 cell-type specific regulatory regions using stratified LD-score regression (S-LDSC)^25^. We found significant enrichments for either of male or female asthma in three annotations; Th0, Th1, and colonic mucosa (*P* < 0.05/220; Supplementary Figure 6). Among them, the colonic mucosa annotation showed significant heterogeneity in the enrichment of heritability (*P*_*het*_ = 0.006 < 0.05/3). Recent studies suggested that host-microbiome interactions at intestinal mucosa (gut-lung axis) have important roles in the development of asthma^26,27^, and our study suggested that the importance of the gut-lung axis in asthma might be different between males and females.

To acquire more insights to disease biology, we estimated the heritability enrichments in the binding sites of a variety of transcription factors (TFs) using S-LDSC. We included TF binding sites defined by 2,868 publicly available chromatin immunoprecipitation sequencing (ChIP-seq) datasets for 410 unique TFs (Supplementary Table 9). To make mutually comparable data, we began our analysis from the raw sequencing data, and defined TF binding sites using a uniform protocol (Methods). Using LD-scores of all TF binding sites, we grouped them into 15 clusters (cluster name was defined by the most dominant TF), and performed uniform manifold approximation and projection (UMAP)^28^ to project all TF binding sites into a two-dimensional space (Methods; Figure 4a and Supplementary Figure 7). To scale the performance of this analysis, we first analyzed previously reported GWAS for red blood cell-related traits^29^ where the critical role of *GATA1* was supported by multiple pieces of evidence^30–34^, and we successfully recapitulated this biology (Figure 4b). We then applied this analysis to our 24 GWAS results (see Methods for selection of diseases), and detected 378 significant enrichments for nine disease (FDR < 0.05) (Figure 4c, Supplementary Figure 8, and Supplementary Table 10). Biologically plausible TFs were highlighted by this analysis; *RELA*, a subunit of NF-κB, for atopic dermatitis, RA, and Graves’ disease; sex hormone receptors (*AR* and *ESR1*) for prostate cancer; and *FOXA2*, which regulates insulin secretion in pancreatic beta-cells^35^, for T2D (Figure 4c). This analysis also suggested that *NKX3-1*, a prostate-specific homeobox gene, has an important role in the biology of prostate cancer (Figure 4c). In addition to this polygenic analysis, the importance of *NKX3-1* was also suggested by the regional analysis integrating eQTL databases; the risk allele of prostate cancer at *NKX3-1* locus (rs4872174-C) was suggested to decrease the expression of *NKX3-1* (Supplementary Table 8). Consistently, loss of *NKX3.1* expression in human prostate cancers was reported to be correlated with tumor progression^36^. Together, our results confirmed and expanded our current understanding of complex traits in the context of TF activity.

In summary, we conducted a large-scale GWAS of 42 diseases in a non-European population and provided rich public resources for genetic studies. Our study provided multiple insights into the etiology of complex traits by integrating annotations of missense variants, eQTL variants, and transcription factor binding site tracks. Currently, genetic studies are overwhelmed by European-descent samples, and the clinical translation of genetic findings would be far more beneficial to European individuals than other populations^1^. Our study contributed to broaden the population diversity in genetic studies and should potentially mitigate the problems originating from this imbalance.

## ONLINE METHODS

### Subjects

All case samples in this GWAS were collected in the BioBank Japan Project (BBJ)^4,5^, which is a biobank that collaboratively collects DNA and serum samples from 12 medical institutions in Japan and recruited approximately 200,000 patients with the diagnosis of at least one of 47 diseases. Among them, cases with dyslipidemia were excluded because it was already reported in our previous study^29^. Amyotrophic lateral sclerosis and febrile seizure were also excluded due to limited sample size. Cases with myocardial infarction, stable angina, and unstable angina were re-classified into a single disease category (coronary artery disease). Thus, we analyzed 42 disease in this study. For control samples, we used samples from the population-based prospective cohorts; the Tohoku University Tohoku Medical Megabank Organization (ToMMo), Iwate Medical University Iwate Tohoku Medical Megabank Organization (IMM)^40^, the Japan Public Health Center–based Prospective Study and the Japan Multi-institutional Collaborative Cohort Study. In addition, we also included samples in BBJ without related diagnoses into control group (Supplementary Figure 1). The sample sizes and the demographic data are provided in Supplementary Table 1. All participating studies obtained informed consent from all participants by following the protocols approved by their institutional ethical committees. We obtained approval from ethics committees of RIKEN Center for Integrative Medical Sciences, and the Institute of Medical Sciences, The University of Tokyo.

### Genotyping

We genotyped samples with the Illumina HumanOmniExpressExome BeadChip or a combination of the Illumina HumanOmniExpress and HumanExome BeadChips. For quality control (QC) of samples, we excluded those with (i) sample call rate < 0.98 and (ii) outliers from East Asian clusters identified by principal component analysis using the genotyped samples and the three major reference populations (Africans, Europeans, and East Asians) in the International HapMap Project^41^. For QC of genotypes, we excluded variants meeting any of the following criteria: (i) call rate < 99%, (ii) *P* value for Hardy Weinberg equilibrium (HWE) < 1.0 × 10^−6^, and (iii) number of heterozygotes less than five. Using 939 samples whose genotypes were also analyzed by whole genome sequencing (WGS), we added additional QC based on the concordance rate between genotyping array and WGS. Variants with a concordance rate < 99.5% or a non-reference discordance rate ≥ 0.5% were excluded. We note that the allele frequency of rs671 (the East Asian-specific functional missense variant at ALDH2) substantially varies among the domestic regions within Japan due to strong selection pressure^42^ and that genotypes of rs671 did not follow HWE. We thus did not apply the HWE QC for rs671. We had confirmed the 100% concordance of rs671 genotypes between the SNP microarray data used in this study and our internal WGS data (n = 2,798; see details in Matoba N. et. al. manuscript in revision).

### Imputation

We utilized all samples in the 1000 Genomes Project Phase 3 (version 5)^7^ as a reference for imputation. We first prephased the genotypes with SHAPEIT2 (v2.778) and then imputed dosages with minimac3 (v2.0.1). After imputation, we excluded variants with imputation quality of Rsq < 0.7. For X chromosome, we performed prephasing and imputation separately for males and females, and we excluded variants with imputation quality of Rsq < 0.7 in either of them.

### Genome-wide association analysis

We conducted a GWAS by employing a generalized linear mixed model (GLMM) using SAIGE (v0.29.4.2)^6^. This strategy enabled us to maintain related samples in our GWAS, and the sample sizes were increased by 6% on average compared to removing related samples. Briefly, there are two steps in SAIGE. In step 1, we fit a null logistic mixed model using genotype data, and we added covariates in this step (see below). In step 2, we performed the single-variant association tests using imputed variant dosages. We applied the leave-one-chromosome-out (LOCO) approach. For the X chromosome, we conducted GWAS separately for males and females, and merged their results by inverse-variance fixed-effect meta-analysis. We used only female control samples for GWAS of female-specific diseases; breast cancer, cervical cancer, endometrial cancer, ovarian cancer, endometriosis, and uterine fibroids. Similarly, we used only male control samples for GWAS of prostate cancer. We incorporated age and top 5 principal component (PC) as covariates. We also used sex as covariate for GWAS of diseases which include both of male and female samples. We created regional association plots by LocusZoom (v1.2)^43^.

We performed stepwise conditional analysis within ± 1 Mb from the lead variant; we repeated the association test by additionally incorporated the dosages of the identified variants as covariates in SAIGE step 1 until we do not detect any significant associations.

We also conducted male-specific and female-specific GWAS using the same pipeline as described above, and estimated heterogeneity in the effect size estimates using Cochran’s Q test.

We set a genome-wide significance threshold at *P* = 5.0 × 10^−8^. For each disease, we defined a significantly associated locus as a genomic region within ± 1 Mb from the lead variant. When a locus did not include any variants which were previously reported to be significantly associated with the same disease (*P* < 5.0 × 10^−8^), we defined it as a novel locus.

### Estimation of heritability

We estimated heritability and confounding bias in our GWAS results with LDSC (v1.0.0)^8^ using the baselineLD model (v2.1)^10^ which include 86 annotations, including 10 MAF- and 6 LD-related annotations that correct for bias in heritability estimates^9^, and were calculated using 481 East Asian samples in 1KG Phase3. For the analysis using LDSC, we excluded variants in the HLA region (chr6:26 Mb-34 Mb). We also calculated heritability Z-score to assess the reliability of heritability estimation.

Absolute quantification of heritability estimation using GWAS results using GLMM can be biased because effective sample size could be different from the true sample size (relative quantification is not biased, and hence GWAS results using GLMM can be applied for genetic correlation analysis and S-LDSC safely). Therefore, to confirm the robustness of heritability estimation in our analysis, we also performed GWAS using generalized linear regression model (GLM). As simple GLM does not account for the bias caused by genetic relationships, we further excluded related samples (Pi-hat by > 0.187), and we analyzed genotype data with PLINK (v1.90)^44^ using the same covariates as described above. Heritability estimates based on GWAS using two different methods (SAIGE vs PLINK) were comparable level (Supplementary Table 2).

### Comparison of GWAS results between populations

To compare the GWAS results of our study with those conducted in European populations, we prepared publicly available GWAS summary statistics for 10 diseases. Summary statistics for eight diseases were downloaded from GWAS Catalog (URL) and their names and their PMID were as follows; atrial fibrillation (30061737), breast cancer (29059683), coronary artery disease (29212778), glaucoma (29891935), ischemic stroke (29531354), prostate cancer (29892016), rheumatoid arthritis (24390342), and type 2 diabetes (30054458). Summary statistics of two diseases were downloaded from UK Biobank GWAS summary statistics at Neale Lab (URL) and their names and their phenotype code were as follows; asthma (22127), and congestive heart failure (I50).

### Pleiotropy

We utilized the following variants detected in GWAS for each disease; (i) lead variants in the significantly associated loci, (ii) independent signals detected by conditional analysis, and (iii) lead variants detected in sex-specific GWAS. We defined pleiotropic association when these variants were in LD (r^2^ > 0.6). We calculated r^2^ using East Asian samples in the 1KG Phase3^7^ by PLINK (v1.90)^44^.

### Functional annotation of associated variants

We utilized the same disease-associated variants as used in the previous section for this analysis. We calculated r^2^ using East Asian samples (r^2^ _EAS_) and European samples (r^2^ _EUR_) in the 1KG Phase3^7^ by PLINK (v1.90)^44^. We annotated disease-associated variants with eQTLs detected in the Japanese population^21^ in the following conditions; (i) the lead variants in eQTL study is in LD (r^2^ _EAS_> 0.6) with GWAS variants and (ii) Q value of the lead variants in eQTL study is less than 0.05. We annotated GWAS variants with eQTL detected in the European population (release v7 of GTEx project)^39^ in the following conditions; (i) the lead variants of eQTL study is in LD (r^2^ _EAS_ > 0.6 and r^2^ _EUR_> 0.6) with GWAS variants and (ii) Q value of the lead variants in eQTL study is less than 0.05.

For the annotation of exonic nonsynonymous variants, we used ANNOVAr^45^. We annotated GWAS variants with nonsynonymous variants when they are in LD (r^2^ _EAS_> 0.6). GRCh37 coordinates were used in this study.

### Genetic correlations between sex-specific GWAS

We estimated genetic correlations between our GWAS results by LDSC (v1.0.0)^8^ using East Asian LD scores which we presented in our previous study^29^. We excluded variants in the HLA region (chr6:26 Mb-34 Mb). We analyzed 20 diseases based on two criteria; (i) heritability was reliably estimated (heritability Z-score > 2; Supplementary Table 2); and (ii) both of male and female patients were included.

### Transcription factor binding sites

We obtained 3,158 raw human ChIP-seq data files in SRA format from the GEO database. We converted them to FASTQ format using the fastq-dump function of SRA Toolkit. We performed QC of sequence reads using FastQC. We mapped these reads to the genome assembly GRCh37 using Bowtie2 (v2.2.5) with default parameters. We called peaks using MACS (v2.1) with default parameters (q < 0.01) and defined them as TF binding sites. We excluded TF binding site tracks which do not have at least one binding region in every chromosome, and 2,868 genome-wide TF binding site tracks remained (Supplementary Table 9).

### Stratified LD score regression

We conducted stratified LD score regression (S-LDSC)^25^ to partition heritability. For S-LDSC analysis of sex-specific GWAS of asthma, we used 220 cell-type specific annotations used in previous articles^25,29^. For other S-LDSC analysis, we used TF binding site tracks which were described in the previous paragraph. For all sites of TF binding, we empirically extended sites by 500 bp at the both ends for this analysis. We computed annotation-specific LD scores using the 1000 Genomes Project Phase 3 (version 5) East Asian reference haplotypes^7^. We estimated heritability enrichment of binding sites of each TF, while we controlled for the merged binding sites of all TFs and the 53 categories of the full baseline model available at the authors’ website (see URLs). We did not use the baselineLD model (v2.1)^10^ in this analysis to avoid false negative findings. We excluded variants in the HLA region (chr6:26 Mb-34 Mb). We analyzed 24 diseases whose heritability was reliably estimated (heritability Z-score > 2; Supplementary Table 2). We calculated the *P* value of the regression coefficient. For each trait, we calculate FDR using the Benjamini-Hochberg method. We set a significance threshold at FDR < 0.05 for this analysis.

### Visualization of TF binding sites

There is a complex correlation structure among 2,868 TF binding site tracks used for S-LDSC analysis. In S-LDSC, we regress GWAS chi-squared statistics on LD-scores of each TF binding site (TF LD-score), and hence we focused on correlations between TF LD-scores, not correlations between TF binding sites. We first performed PCA using all TF LD-scores. To classify them into mutually correlated TF groups, we performed k-means clustering (k=15) using top 15 PCs. We named each cluster by the most dominant TF in each cluster (Figure 4). The list of each TF binding site and its assigned cluster name was provided in Supplementary Table 9. We then performed uniform manifold approximation and projection (UMAP)^28^ using top 15 PCs to project all TF binding sites into a two-dimensional space. UMAP was conducted using R package umap (v.0.2.0.0). Our workflow was illustrated in Supplementary Figure 7.

### Data availability

GWAS summary statistics of the 40 diseases (all except breast cancer and coronary artery disease) are publicly available at our website (JENGER; see URLs) and the National Bioscience Database Center (NBDC) Human Database (Research ID: hum0014) without any access restrictions. For breast cancer and coronary artery disease, we will deposit the results after acceptance to the journal due to accompanying projects. GWAS genotype data for case samples were deposited at the NBDC Human Database (Research ID: hum0014).

## URLs

BBJ, https://biobankjp.org/english/index.html

JENGER http://jenger.riken.jp/en/

PLINK 1.9, https://www.cog-genomics.org/plink2

MACH, http://csg.sph.umich.edu//abecasis/MaCH/

Minimac, https://genome.sph.umich.edu/wiki/Minimac

SAIGE, https://github.com/weizhouUMICH/SAIGE

GWAS Catalog, https://www.ebi.ac.uk/gwas/

Neale Lab, http://www.nealelab.is/uk-biobank

PASCAL, https://www2.unil.ch/cbg/index.php?title=Pascal

ldsc, https://github.com/bulik/ldsc/

LD score, https://data.broadinstitute.org/alkesgroup/LDSCORE/

ANNOVAR, http://annovar.openbioinformatics.org/en/latest/

Locuszoom, http://locuszoom.sph.umich.edu/locuszoom/

R, https://www.r-project.org/

SRA Toolkit, https://www.ncbi.nlm.nih.gov/sra/docs/toolkitsoft/

FASTQC, https://www.bioinformatics.babraham.ac.uk/projects/fastqc/

Bowtie2, http://bowtie-bio.sourceforge.net/bowtie2/manual.shtml

MACS, https://github.com/taoliu/MACS

NBDC Human Database, https://humandbs.biosciencedbc.jp/en/.

1000 Genomes Project, www.1000genomes.org/

## ACKNOWLEDGMENTS

We acknowledge the staff of BBJ for their outstanding assistance. We express our heartfelt gratitude to Tohoku University Tohoku Medical Megabank Organization (ToMMo), Iwate Medical University Iwate Tohoku Medical Megabank Organization (IMM), the Japan Public Health Center–based Prospective (JPHC) Study, and the Japan Multi-Institutional Collaborative Cohort (J-MICC) Study for their invaluable contributions to collecting control samples. We also express our gratitude to E.K. and H.S. for kindly sharing their results of ChIP-seq data analysis. We extend our appreciation to Y. Yukawa, Y. Yokoyama, and other members of the Laboratory for Statistical Analysis, RIKEN Center for Integrative Medical Sciences for their great support. This research was supported by the Tailor-Made Medical Treatment Program (the BioBank Japan Project) of the Ministry of Education, Culture, Sports, Science, and Technology (MEXT) and the Japan Agency for Medical Research and Development (AMED). The JPHC Study has been supported by the National Cancer Center Research and Development Fund since 2011 and was supported by a Grant-in-Aid for Cancer Research from the Ministry of Health, Labour and Welfare of Japan from 1989 to 2010. The study of psychiatric disorders was supported by AMED under Grant Numbers JP18dm0107097, JP18km0405201 and JP18km0405208.

## AUTHOR CONTRIBUTIONS

K.Ishigaki wrote the manuscript with critical inputs from S.R and Y.Kamatani. K.Ishigaki conducted all bioinformatics analyses with the help of M.A, M.Kanai, A.T, S.S, N.Matoba, S.K.L, Y.O, C.Terao, T.A, S.G, S.R, and Y.Kamatani. Y.Momozawa and M.Kubo performed genotyping. H.S and E.K. analyzed ChIP-seq data. M. Ikeda and N. I managed GWAS data of psychiatric diseases. S.K.L, Y.Kochi, M.Horikoshi, Ken Suzuki, K.Ito, M.Hirata, K.M, S.I, I.K, T.Tanaka, H.N, A.Suzuki, T.H, M.T, K.C, D.M, M.M, S.N, Y.D, Y.Miki, T.Katagiri, O.O, W.O, H.I, T.Yoshida, I.I, T.Takahashi, C.Tanikawa, T.S, N.Sinozaki, S.Minami, H.Yamaguchi, S.A, Y.T, K.Yamaji, K.T, T.F, R.T, H.Yanai, A.M, Y.Koretsune, H.K, M.H, S.Murayama, K.Yamamoto, Y.Murakami, Y.N, J.I, T.Yamauchi, T.Kadowaki, M.Kubo, and Y.Kamatani contributed to the management of BBJ data. N.Minegishi, Kichiya Suzuki, K.Tanno, A.Shimizu, T.Yamaji, M.Iwasaki, N.Sawada, H.U, K.Tanaka, M.N, M.S, K.W, S.T, and M.Y contributed to the management of cohort control data. S.R, J.I, T.Yamauchi, T.Kadowaki, M.Kubo, and Y.Kamatani jointly supervised this study.

