## Supplementary Figures for "Large scale genome-wide association study in a Japanese population identified 45 novel susceptibility loci for 22 diseases"

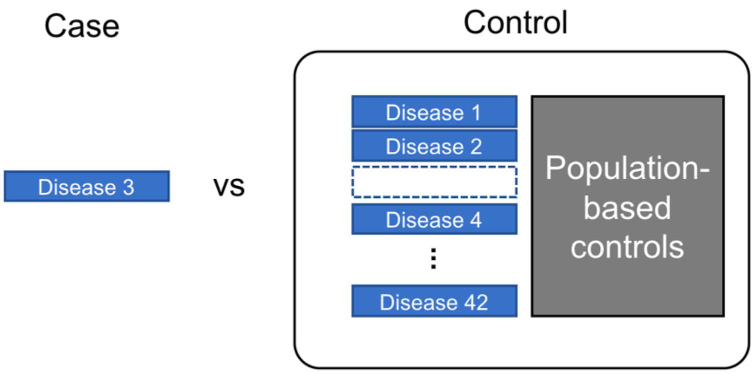


**Supplementary Figure 1. Study design of this GWAS.**

An example of our study design. In GWAS of disease 3, we included all other patients (except those have related diseases) into control group. For example, for GWAS of asthma, patients with COPD were excluded from control samples. The definition of related diseases is provided in Supplementary Table 1.

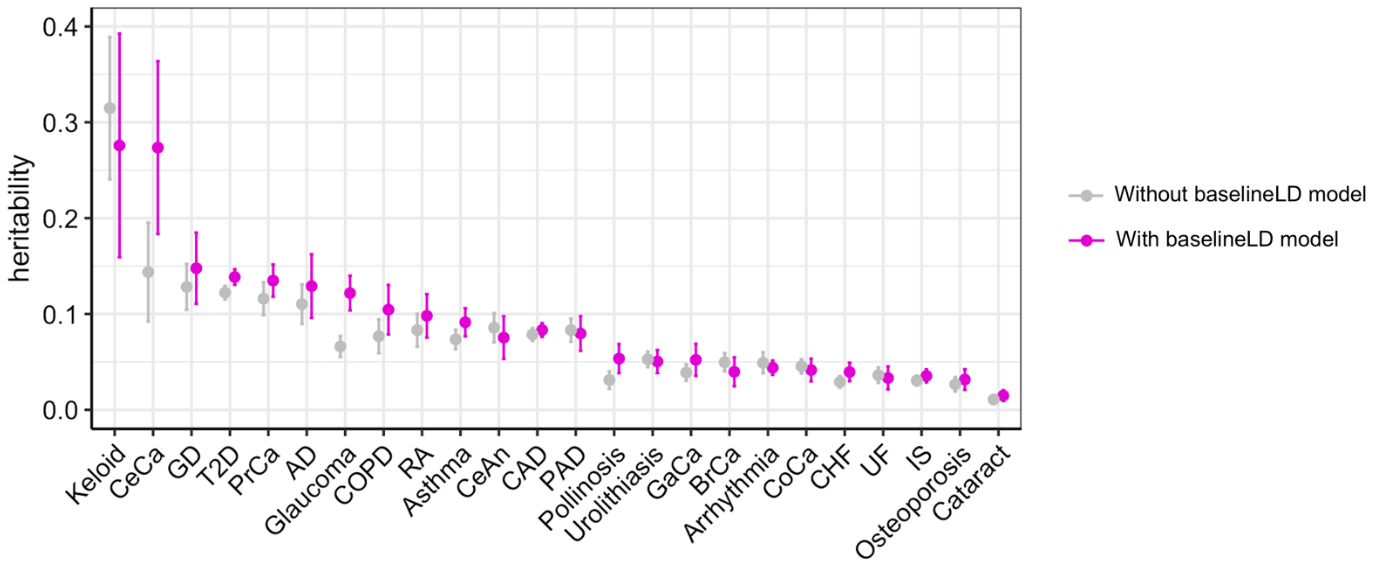


**Supplementary Figure 2. BaselineLD model improved heritability estimation using LDSC.**

The estimates of liability scale heritability were shown. Heritability was estimated by LDSC with or without using baselineLD model v2.1. GWAS summary statistics of SAIGE were used for this plot.

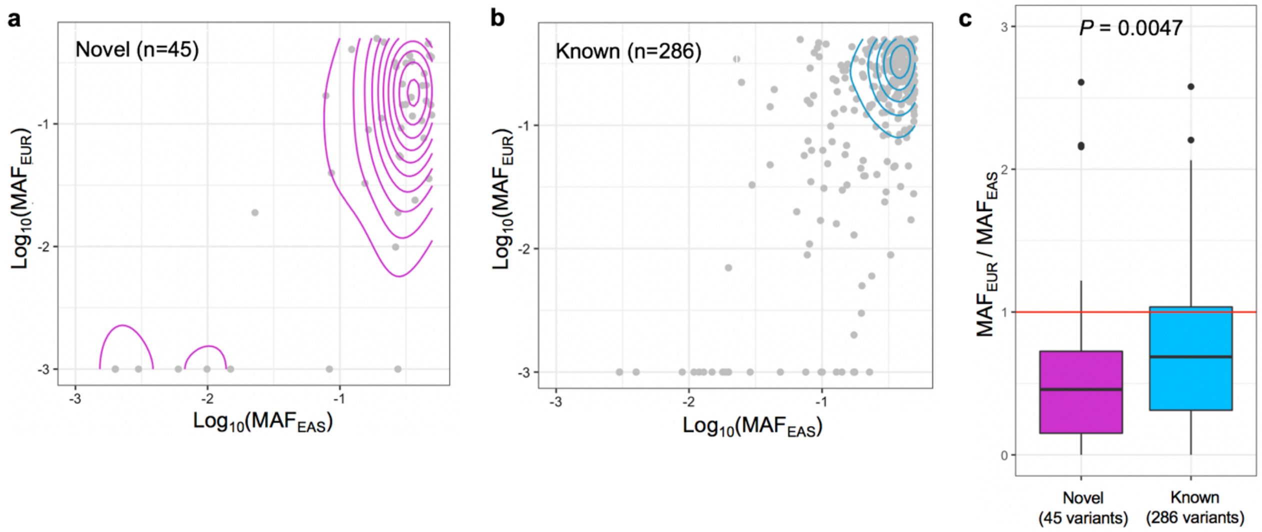


**Supplementary Figure 3. Comparison of allele frequencies at disease-associated variants between East Asian and European populations.**

**a and b**, Allele frequencies in 1KG Phase3 at the novel and known variants were plotted. When MAF < 0.001, MAF was adjusted to 0.001 to fit in log scale. $MAF_{EAS}$, MAF in East Asian populations (1KG Phase3). $MAF_{EUR}$, MAF in European populations (1KG Phase3). **c,** The ratios of allele frequency ($MAF_{EUR}/MAF_{EAS}$) were provided. The differences in ratios were tested by Mann–Whitney U test, and its *P* value was provided.

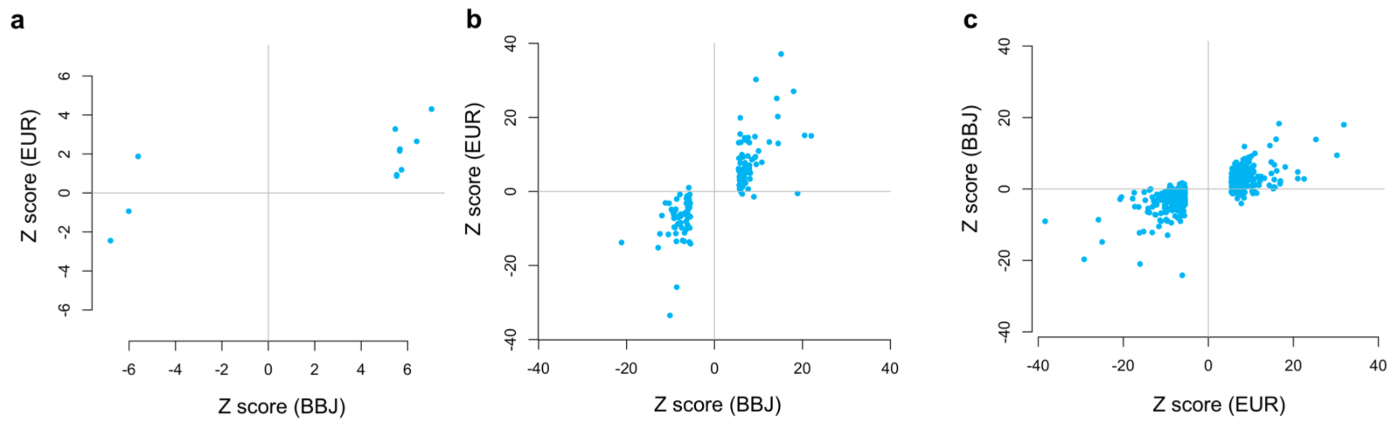


**Supplemental Figure 4. Comparison of allelic directions between our GWAS and previous European GWAS. a,** statistics of the lead variants at the novel loci in our study (n = 11). **b,** statistics of the lead variants at the known loci in our study (n = 146). **c,** statistics of the lead variants at the significant loci in European studies (n = 665). Z score (BBJ), Z scores of the alternative allele in our study. Z score (EUR), Z scores of the alternative allele in European GWAS.

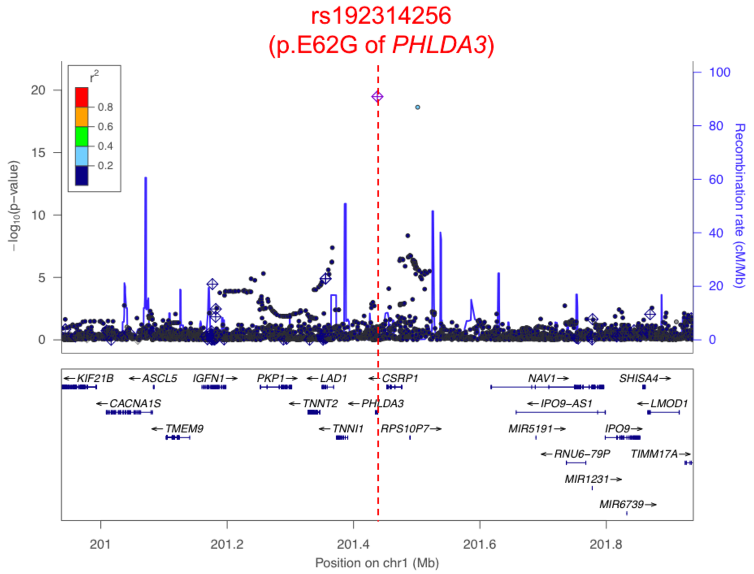


**Supplementary Figure 5. A novel association which can be explained by an East Asian-specific missense variant.**

A regional association plot for keloid at the *PHLDA3* region is provided.

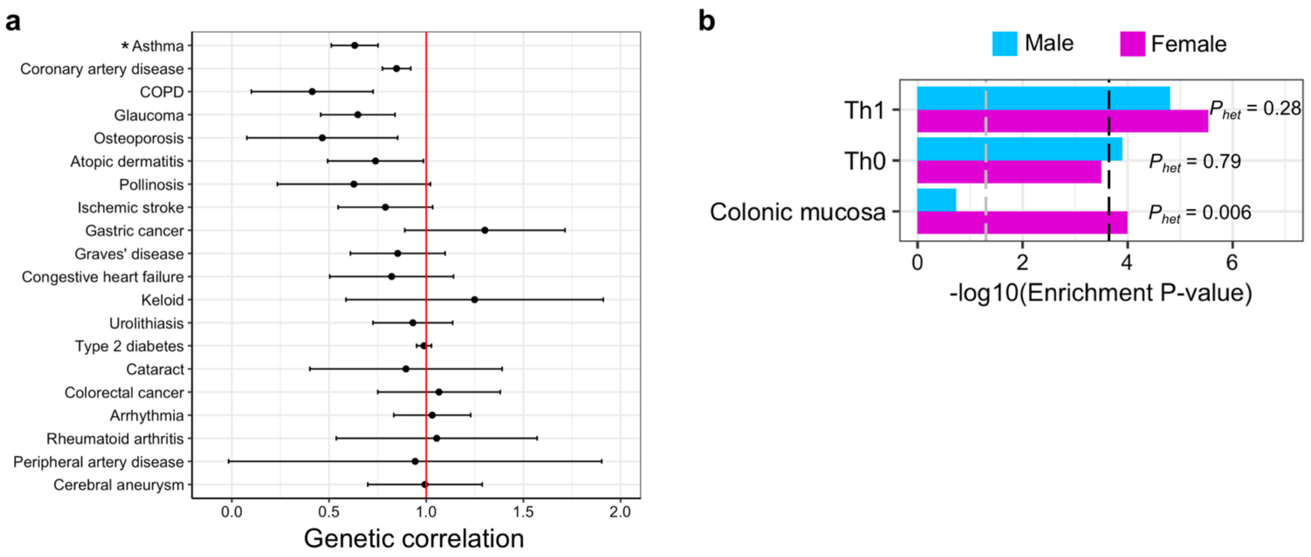


**Supplemental Figure 6. Genetic correlations between male- and female-specific GWAS.**

**a.** Genetic correlations between male- and female-specific GWAS. Estimates of genetic correlation and standard errors are provided. *: genetic correlation was significantly different from one ($P$ < 0.05/20). **b.** The results of S-LDSC analysis based on sex-specific GWAS of asthma using 220 cell-type specific annotations. Significant annotations in either male or female asthma were shown ($P$ < 0.05/220). Heterogeneity was tested by Cochran's Q test, and its $P$ values ($P_{het}$) were also provided. Black dashed line indicates $P$ value = 0.05/220; grey dashed line indicates $P$ value = 0.05.

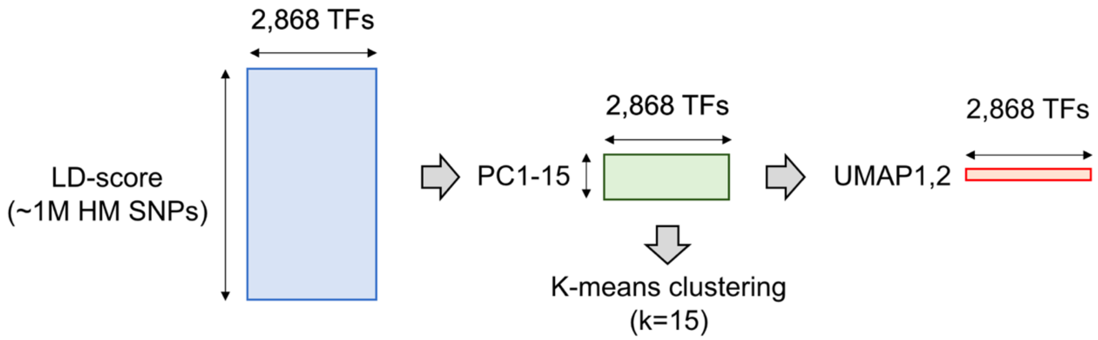


**Supplemental Figure 7. Our workflow to visualize TF annotations in two-dimensional space.**
We first performed PCA using LD-score of 2,868 TF binding sites. To classify them into mutually correlated TF groups, we performed k-means clustering (k=15) using top 15 PCs. We named each cluster by the most dominant TF in each cluster (Figure 3). We then performed uniform manifold approximation and projection (UMAP) using top 15 PCs to project all TF binding sites into a two-dimensional space. A rectangle indicates a matrix.

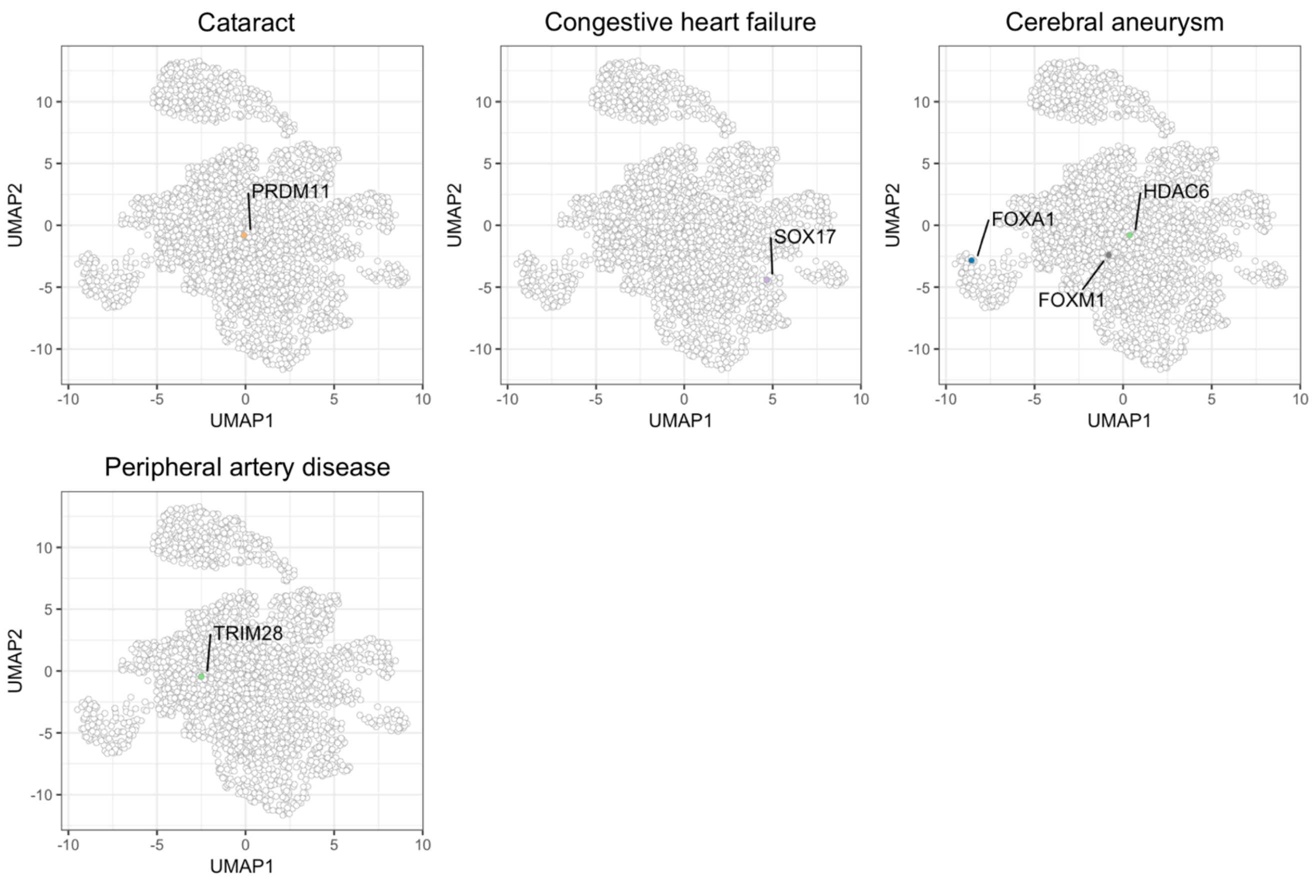


**Supplemental Figure 8. S-LDSC results of four diseases in our GWAS.**

The results of S-LDSC were plotted on the UMAP space. The significant results (FDR<0.05) were highlighted by cluster-specific colors (the same colors as used in Figure 4). The names of the top five most significant TFs were also shown on the plot. The results of diseases with less than five significant TF binding site tracks were shown.
